# Fast and robust method for drug response biomarker identification and sample stratification

**DOI:** 10.1101/525337

**Authors:** Simanti Bhattacharya, Amit Das

## Abstract

With unprecedented progress of cancer research, the world is now prepared with versatile arsenal of drugs to combat cancer. However, individual’s response to any drug or combination treatment stands as a major challenge and hence there exists the sheer need for personalized medication. Identification of drug response biomarkers from a wholistic tumor microenvironment analysis would guide researchers to develop custom-tailored treatment regimen.

In this study, a fast and robust method has been developed to identify drug response biomarkers from entire transcriptomics data analysis in a data-driven manner. The biomarkers which were identified by the method, were able to stratify patients between responders vs non-responders population. Furthermore, bayesian network (BN) analysis, done on the data, brought forth a mechanistic insight into the role of identified biomarkers in regulating drug’s efficacy.

The importance of this work lies with the protocol that is time saving and requires less computation power, yet analyzes a whole system data and helps the researchers to take a step forward towards the development of personalized care in effective cancer treatment.

**Highlights:**

1. Supervised machine learning approach to analyze gene expression data.
2. Drug response biomarker identification.
3. Categorization of samples for their drug response with the help of identified biomarkers.
4. Functional enrichment to understand the biomarkers association with biological processes.
5. Bayesian network analysis to develop causal structure among identified biomarkers and drug targets.
6. Time and cost-effective pipeline for fast and robust prediction of drug response biomarkers.

## 1. Introduction

The development of the tumor and its microenvironment is dependent on multilevel systemic regulations [1]. Tumor developmental stages, benign to malignant transition [2], tumor immune escape mechanism [3], epigenetic modulation [4] as well as disease specific mutational switches [5] etc., all regulate the disease prognosis. In addition to these intrinsic modulators, external risk exposure, such as smoking habit, life style, eating habit etc adds another level of complexity [6,7]. Interestingly, with the unmatched speed of cutting-edge technologies and researches, present world is much prepared with essential therapies and medications [8–11], monotherapy or combination, to satisfy the need (1) to fight against cancer progression, (2) to increase the patient’s life expectancy and (3) to prohibit the relapse of the disease. Effective cancer therapies can be of several types [12]: surgery (surgical removal of affected tissue), radiation (high doses of radiation applied to kill cancer cells and shrink tumors), chemotherapy (use of chemotherapeutic agents to kill cancer cells), immune therapy (to elicit effective immune system to inhibit tumor cell’s immune escape and to destroy tumor cells precisely) and targeted therapy (use of specific drugs that targets cancer specific antigens, receptors etc). For certain cancers, hormone therapy and stem cell transplantation have shown very promising results. These medications, as monotherapy or as a combination, have been very effective to pull the chain of rapid spread of cancer.

However, there exists a gridlock that obstructs these cancer therapies to be used effectively in an individual patient. Owing to the complexity of tumor microenvironment, an individual patient responds differently to the given treatment regimen. Hence arises the need for personalized medication [13–15] wherein the treatment regimen will be custom-tailored for the patient to have the best outcome of the therapy. In order to fulfill this efficient treatment, the identification of drug response biomarkers [16] becomes essential. Identification of drug response biomarkers has become an integral part of modern-day personalized therapy [17,18]. With the advancement of data-driven approaches [19,20], the concept behind ‘one-drug-one-target’ has changed. The response outcome of a drug depends on many factors, such as mutations, compensatory mechanisms, epigenetics, dysregulated genes and many more, making a prediction of drug’s efficacy a real challenge for decades. These regulatory factors either could be in the immediate vicinity of drug’s mechanism of action (MoA) or could be trans-regulatory that acts by modulating relevant genes’ expression. These down-streams effects (i.e. gene’s up or down-regulations) are easy to assess and can directly be linked to pathways or cellular processes, thereby, giving an answer to how and why a patient could be sensitive to a given therapy [21]. The advantage of such a data-driven approach is that it is free from any pre-conceptualized bias like drug targets, disease genes etc.

However, dealing with the enormous amount of gene expression data is both time and computation expensive. There is a sheer need for a rapid and robust method for biomarker identification which can lead to stratify patients as per their response to the drug. In this study, transcriptomic data has been analyzed through a machine learning pipeline which has been developed for rapid identification of drug response biomarkers and thereafter patient stratification.

The work has used the data from the experiment submitted to NCBI GEO as GSE2535 [22] where chronic myeloid leukemia (CML) is the disease system and imatinib is the drug. Briefly, CML is a myeloproliferative disorder caused by constitutive tyrosine kinase activity of Bcr-Abl oncoprotein. Although Bcr-Abl-inhibitor imatinib stands the first-line therapy against CML, almost 20-30% of patients develop drug resistance [23]. In general, mutations in the Bcr-Abl domain increase the formation of imatinib resistance. However, not all resistant patients are Bcr-Abl mutation dependent which triggers the thought that there exist non-mutant mechanisms leading to imatinib resistance.

After analyzing the transcriptomic data, 12 genes have come up as potential biomarkers which have efficiently categorized unlinked patient samples by their drug response. Further to this analysis, potential links between this drug and the identified biomarkers have been established through function enrichment. However, identification of biomarkers, coupled with their functional analysis may not satisfy the query regarding the causality behind these potential biomarkers and drug’s activity. To understand the system-level regulatory mechanism, here, bayesian network (BN) analysis [24,25] has been employed which developed a probabilistic model of the network with structural characteristics and directed acyclic networks in two different classes: responders vs non-responders so as to enlighten the regulatory map behind the scene.

## 2. Materials and methods

### 2.1. Patients description

Patient description was explicitly described in [22]. Authors collected two sets of patients: (1) chronic myeloid leukemia (CML) patients at chronic phase 1 from Novartis-sponsored trials at the Department of Hematology of the University of Leipzig, Germany (n=15) and (2) German CML Study Group at the Faculty of Clinical Medicine Mannheim of the University of Heidelberg, Germany (n=14). In both cases, the patient’s response to imatinib was captured and cellular lysates for RNA extraction were stored prior to imatinib therapy. Later on, one patient’s data was ignored due to defective chip data. Hence total 28 patient samples were analyzed in the work (GSE2535; [22]) which was also utilized in current research. There was no significance difference in average age as well as the duration of the disease between two sets of patients. There were 15 female and 13 male patients.

Responders and non-responders were defined as follow: Responders (R) to imatinib who had achieved complete cytogenetic response (CCR) <9 months (n=16), while non-responders (NR) to imatinib who had failed to achieve a major cytogenetic response (MCR) within one year of treatment (n=12).

### 2.2. RNA extraction and microarray

Details of the sample collection methods and equipment were explained in [22]. Briefly, bone marrow or peripheral blood samples were collected from these patients, before Imatinib treatment. RNA was extracted either with RNeasy kit or cesium-chloride gradient purification method. Quality of RNA extracted was assured with the presence of discrete 18S and 28S ribosomal RNA peaks and the absence of irregularly-sized low molecular weight RNA species in the electropherogram. The microarray was performed on Affymetrix chip following standard protocol. Raw samples were downloaded from NCBI GEO and then were normalized with GCRMA.

### 2.3. Data processing

The normalized matrix had 12625 genes. Median over genes were done to obtain 9204 unique genes. Samples were labeled according to the response (coded as “1”) o resistance (coded as “-1”) towards the drug. Samples were then randomly shuffled.

### 2.4. Machine learning

#### 2.4.1. Model building

20 samples were used to build the model and to train it. Remaining 8 samples were kept aside from the beginning that would be used as an external or unlined test sample to validate the output. Details of the workflow have been given in Fig. 1.

**Fig. 1.**
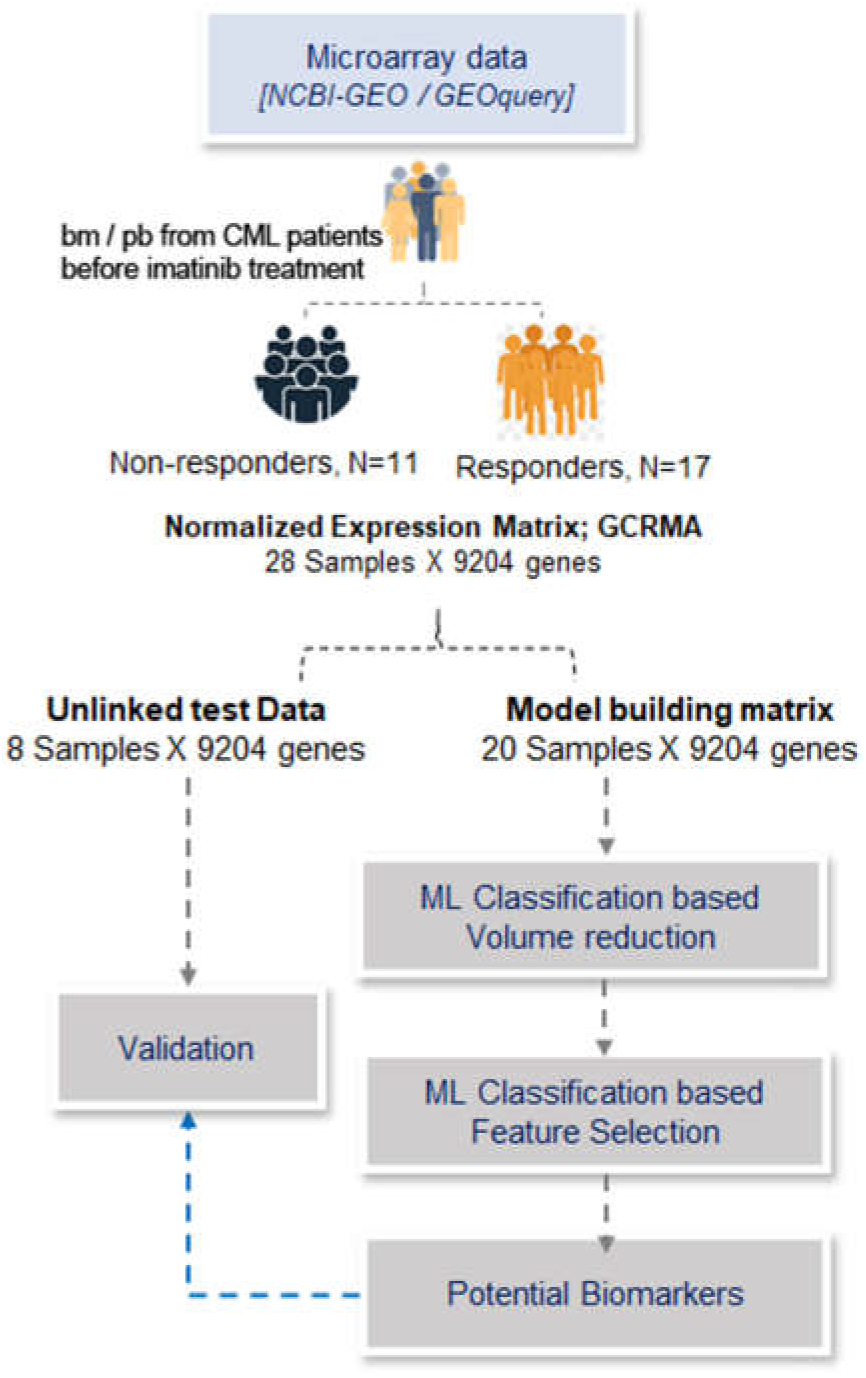
Detailed workflow for biomarker selection. This is a schematic diagram of the pipeline used in this study. The work was started with a microarray data 9204 genes and 28 CML patients whose drug response was known. Bone marrow (BM) or Peripheral blood (pb) samples for microarray analysis were collected before imatinib treatment. For the main model building, 20 samples were used while 8 samples were kept aside for validation. Different ML algorithms were used stepwise to obtain drug response genes.

The matrix had samples in rows and genes in columns. These genes were considered as variables and genes with the expression pattern were considered as features. In the machine learning algorithm, this 20 samples, were further subdivided into training samples (75% of 20 samples) and testing samples (25% of 20 samples). Parameters were optimized to improve the prediction of this test set response.

#### 2.4.2. Feature selection

Feature selection step was preceded by volume reduction with <u>**V**</u>ariable <u>**S**</u>election <u>**U**</u>sing <u>**R**</u>andom<u>**F**</u>orest (VSURF [26]) which uses the randomForest method to select significant variables in case of high dimensional data. The reduced matrix had all insignificant gene pruned out and only relevant ones were then given as input in feature selection step where randomForest [27] classification was used. Classification based analysis was done with 500 trees and 34 splitting at each tree node. Out-of-bag error estimation was also calculated. Class error probability is given in the table below Variables were selected based on their importance weight which was equivalent to their contribution towards the sample’s drug response.

#### 2.4.3. Validation

The knowledge of final features was then applied to the 8 samples, kept aside as external test cases to predict their response pattern. Finally, observed and predicted responses were compared to understand the prediction accuracy of the selected feature genes. For validation, 3 different prediction algorithms (<u>**S**</u>upport <u>**V**</u>ector <u>**M**</u>achine: SVM [28], <u>**r**</u>andom<u>**F**</u>orest: RF and <u>**k**</u>-<u>**N**</u>earest <u>**N**</u>eighbor: k-NN [21]) were used to increase the confidence and robustness of prediction.

### 2.5. Enrichment analysis

Gene ontology (GO) [29,30] to gene association data for Homo sapiens was downloaded from gene ontology consortium (*http://www.geneontology.org/*). GO namespace and category were retrieved from .obo file. These two files were used to create GO to gene dataset which was further used to enrich GO terms (only biological processes, BP) for the selected genes. Hypergoemetric p-value was generated to obtain relevant GO-BP terms.

### 2.6. Bayesian network

Bayesian networks (BN) [24,25,31] are a type of probabilistic graphical model that can be used to build models from data and/or expert opinion. BN is also called a directed acyclic graph or DAG. It constitutes of nodes (each node represents a variable such as genes. A variable might be discrete, such as drug response = {Yes, No} or might be continuous such as gene expression values) and links/edges (connections with directions across node pairs). The absence of link between a pair of nodes does not mean that they are completely independent; rather they may be connected via other nodes. However, such node pairs may become dependent or independent depending on the evidence of other nodes. Each gene pair has a source node/gene and a target node/gene which is associated with a strength value (probability of having this gene pair) and a direction value (probability of having the said direction).

In this study, expression data-driven BN was constructed using bnlearn package [32] in R with an aim decipher potential gene regulatory networks or protein interaction networks. Separate networks were created for the responders and the non-responders group. For each group, the input dataset was a matrix with samples in rows and genes in columns. Both datasets had 20 genes (Machine learning derived genes paired with known drug target genes). While the responders dataset had 17 samples, the non-responders dataset had 11 samples. Each BN was validated through k-fold cross validation and post boot-strapping, arcs or edges with a minimum of 75% direction and strength value were selected and compared through the R compareDF package.

## 3. Results and discussions

### 3.1. Feature selection

Biological data is unique in nature where variable size (number of genes) is much greater than sample (number of observations) size. Moreover, variables i.e. genes have an influence on each other either directly or through some other genes [33]. The enormous variable size creates computation issues while performing machine learning on such data in the local system. The high dimension of data must be reduced so that machine learning can be performed on less number but significant genes.

In this work, the entire gene expression matrix (with 9204 genes) was used (Fig. 2A). From this enormous dataset, significant genes were selected using VSURF algorithm [26] which selects important variables by recursive removal of insignificant or low weight genes. Hence, the matrix dimension in our case, reduced to 1184 genes and 20 samples (Fig. 2B). Performing all three steps of VSURF is a time-consuming process. That is why only VSURF threshold step was performed.

**Fig. 2.**
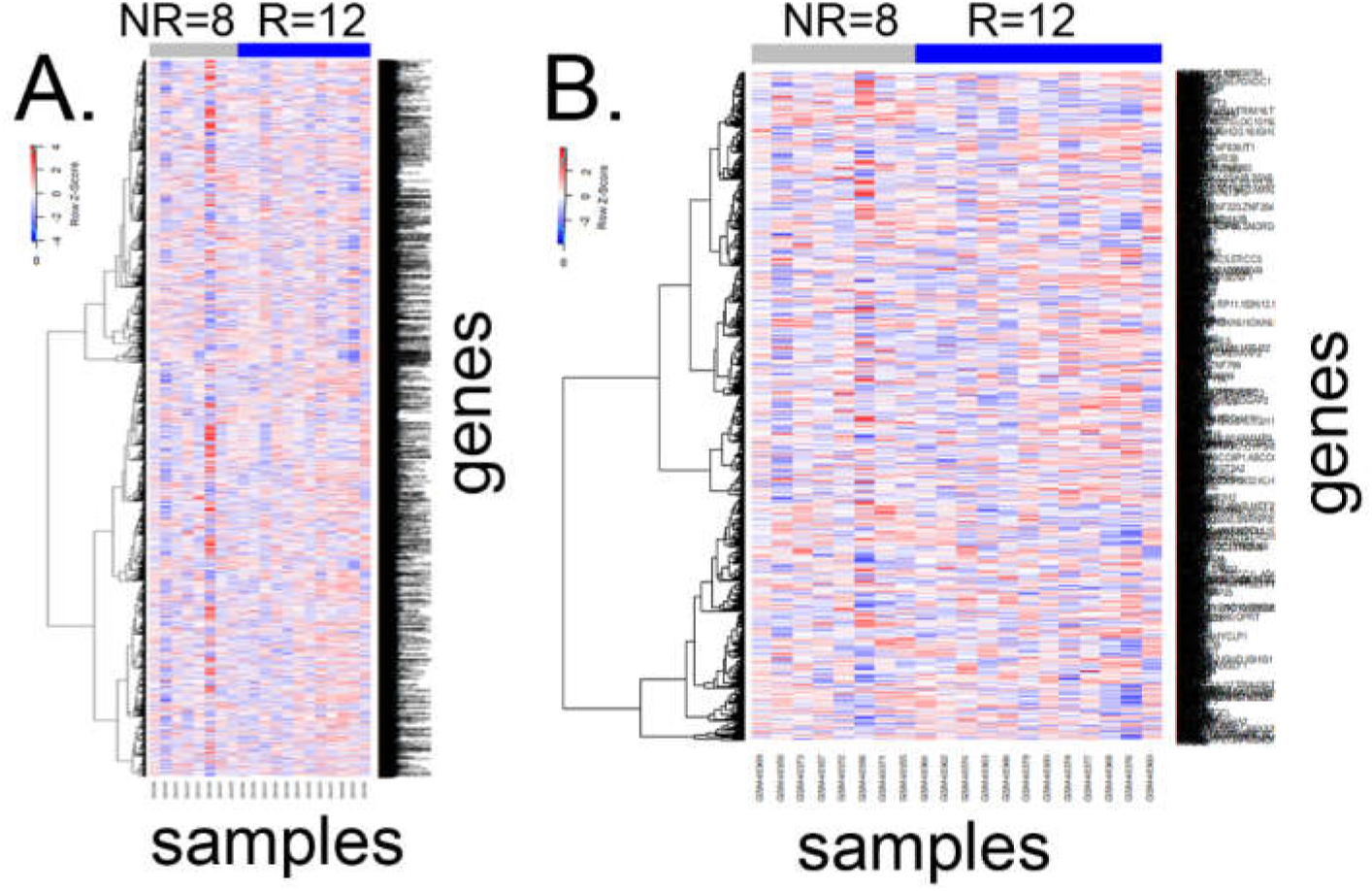
Heatmap for gene expression pattern. Heatmaps for entire gene set, n=9204 (A) and after VSURF mediated volume reduction, n=1184 (B) are shown here. NR stands for non-responders, 8 samples and R stands for responders, 12 samples. This large set of genes are not distinctly different in both classes of samples. In both cases, samples are shown in the x axis and genes are shown in the y axis. During heatmap generation, expression values were scaled. Given color-scale indicates that comparative lower expressions are shown in shades of blue while comparative higher expressions are shown in shades of red.

Thereafter RF was applied to this model system which estimated the importance or weights of each predictor variable (here, gene) in modelling the classes (“1” & “-1”). These weights were equivalent to genes importance in classifying samples into two distinct classes. 40 genes were selected with weight >=0.15 criteria (Fig. 3A) and with subsequent steps finally 12 genes were selected (Fig. 3B) that showed distinct expression pattern between two classes.

**Fig. 3.**
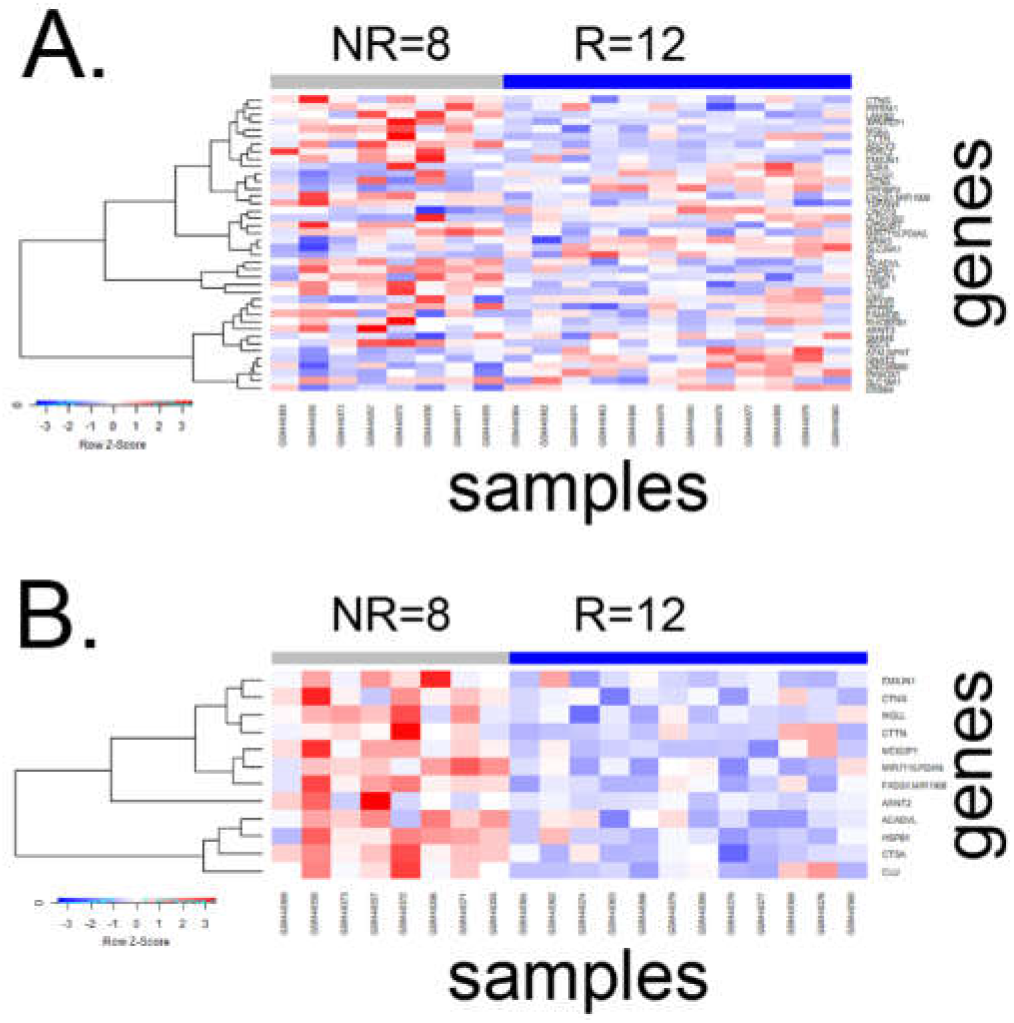
Feature selection using a data-driven approach. (A) With randomForest cut-off >0.15, 40 genes were selected. The heatmap shows there exist patches of difference in genes expression between responders(R) and non-responders(NR). (B) with further shrinking, 12 genes were selected. Their expressions are strikingly different between two classes, i.e. R vs NR. In both cases, again, samples are shown in the x axis and genes are shown in the y axis. During heatmap generation, expression values were scaled. Given color-scale indicates that comparative lower expressions are shown in shades of blue while comparative higher expressions are shown in shades of red.

The identified potential biomarkers with their weights have been given in table 2.

**Table 1.** Class error probability for randomForest prediction. “-1” stands for non-responder and “1” stands for responder class. For non-responder and responder classes, there are 0.25% and 0.08% error probability, respectively for the randomForest training model.

|  |  | *Predicted* |  |  |
| --- | --- | --- | --- | --- |
|  |  | non-responder(-1) | responders (1) | class.error |
| *Observed* | non-responder (-1) | 6 | 2 | 0.25 |
|  | responder (1) | 1 | 11 | 0.083333 |

**Table 2.** Identified biomarkers with their weights. Identified biomarkers (i.e. genes) with their ML weights are given in the table. A negative weight corresponds to comparative lower expression in drug-responding samples. Gene names and Gene IDs are also mentioned.

| GENE SYMBOL | WEIGHT | GENE NAME | GENE ID |
| --- | --- | --- | --- |
| CLU | -0.5344 | Clusterin | 1191 |
| HSPB1 | -0.2965 | Heat shock protein family B small member 1 | 3315 |
| ARNT2 | -0.25353 | Aryl hydrocarbon receptor nuclear translocator 2 | 9915 |
| MGLL | -0.24054 | Monoglyceride lipase | 11343 |
| **CTSA** | -0.21896 | Cathepsin A | 5476 |
| **CTTN** | -0.20909 | Cortactin | 2017 |
| **MIR7110.PDIA5** | -0.20887 | Protein disulfide isomerase family A member 5 | 10954 |
| **ACADVL** | -0.18557 | Acyl-CoA dehydrogenase very long chain | 37 |
| **EMILIN1** | -0.16605 | Elastin microfibril interfacer 1 | 11117 |
| **CTNS** | -0.16163 | Cystinosis, lysosomal cystine transporter | 1497 |
| **FADS1.MIR1908** | -0.16033 | Fatty acid desaturase 1 | 3992 |
| **MEIS3P1** | -0.15428 | Meis homeobox 3 pseudogene 1 | 4213 |

### 3.2. Feature selection-based sample stratification

A sub matrix was then formed with these 12 genes and 20 samples which was further used to train model of three different algorithms: SVM, RF and k-NN. All these three algorithms are distinct from each other. This trained model was used to predict the response of external test data (8 samples) with 75% accuracy (table 3).

**Table 3.** Feature selection-based sample stratification. The identified biomarkers and their weights were used to determine the response outcome of remaining 8 samples, unlinked to the main training data. Three different prediction algorithms were used to predict the outcome. The “Inference” column in the table, holds the cumulative prediction from all three prediction algorithms. A comparison between “Inference” and “Actual Response” declares that the pipeline has predicted the outcome of 6 out of 8 samples correctly, i.e. 75% accuracy.

|  |  | Different predictive algorithms |  |  |  |
| --- | --- | --- | --- | --- | --- |
| Unlinked Sample N=8 | Actual Response | k-NN | randomForest | SVM | Inference |
| GSM48381 | Responder | Non-Responder | Responder | Responder | Responder |
| GSM48370 | Non-Responder | Non-Responder | Non-Responder | Non-Responder | Non-Responder |
| GSM48354 | Non-Responder | Responder | Responder | Responder | Responder |
| GSM48361 | Responder | Responder | Responder | Non-Responder | Responder |
| GSM48375 | Responder | Responder | Responder | Responder | Responder |
| GSM48365 | Responder | Responder | Responder | Responder | Responder |
| GSM48367 | Non-Responder | Responder | Responder | Responder | Responder |
| GSM48359 | Responder | Responder | Responder | Responder | Responder |

### 3.3. Functional analysis

Identified genes that could predict the response of samples to the drug, could be used as potential drug response biomarkers. Hence it was required to understand their link to drug’s mechanism of action. Most often the not, it has been found that the identified genes might regulate the system distantly, instead of direct regulation, culminating to alternation in response to the drug. In order to decipher the processes, GO functional enrichment was carried out with the selected genes. This analysis enriched 70 terms (FDR<0.05). These terms were very specific which were required to be clustered to have a broad category. Revigo webtool (*http://revigo.irb.hr/*) was used to categorize the specific GO terms into six distinct classes and one generic class, grouped as ‘other’ (Fig. 4).

**Fig. 4.**
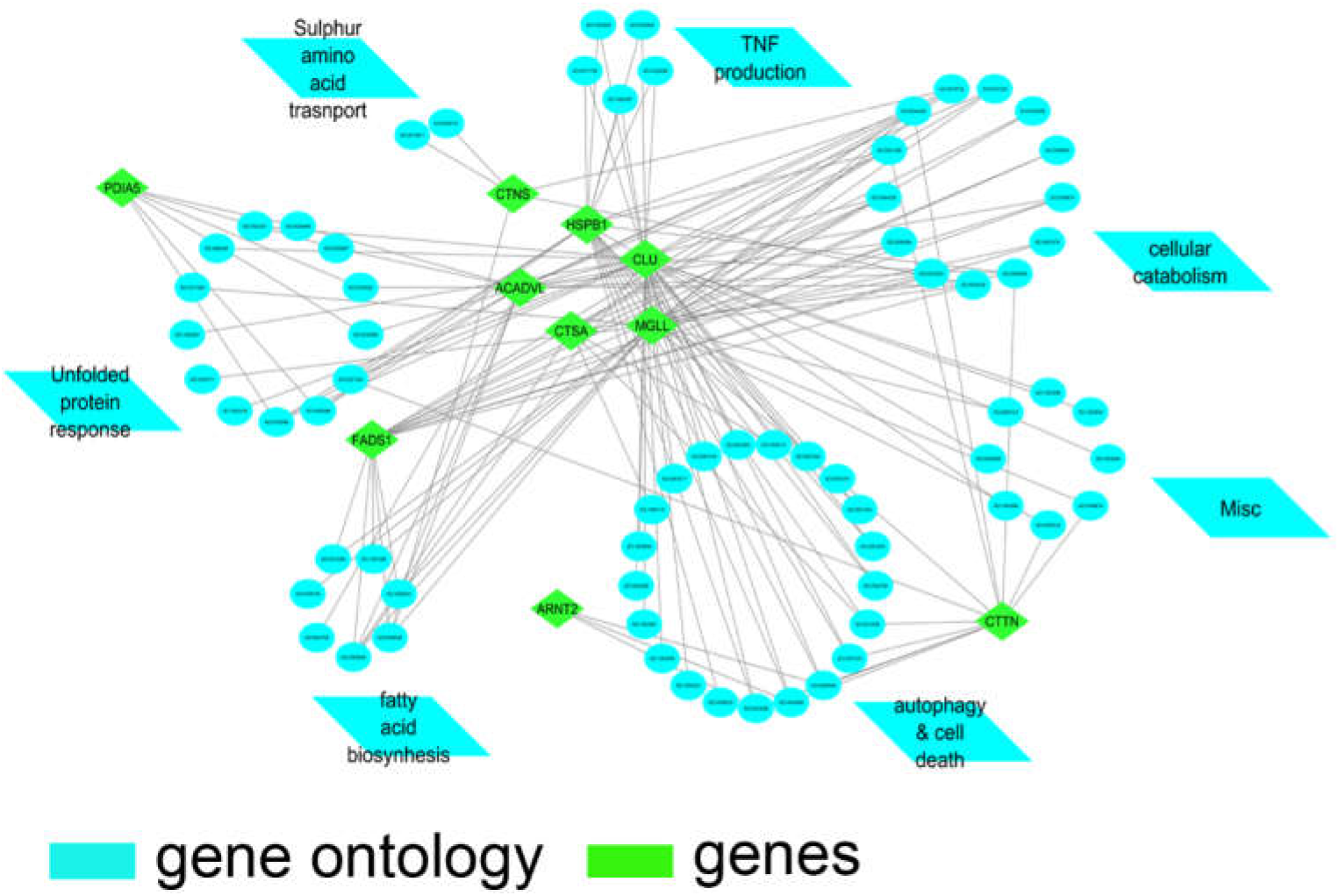
Functional enrichment analysis. Gene ontology enrichment was carried out with hypergeometric p-value calculation using identified biomarkers or feature genes (green, rhombus). Selected significant gene ontology (cyan, circle) terms (FDR <0.05) were further categorized into broad clusters (cyan, parallelogram) using Revigo webtool.

Published experiments were reviewed to establish the link between drug’s efficacy and the enriched processes.

#### 3.3.1. Regulation of autophagy

Autophagy enables recycling of intracellular ingredients as an alternative source during metabolic stress or starvation, especially in cancer cells, to maintain cellular homeostasis and survival. Helgason *et al*.,(2011) experimentally proved that autophagy inhibition in combination with TKIs (imatinib, dasatinib, or nilotinib) resulted in almost elimination of CML stem cells [34]. Moreover, overexpression and knockdown of WNT2 have demonstrated perturbations in the autophagy process and thereby CML drug sensitivity to Imatinib.

#### 3.3.2. Response to topologically incorrect protein

Topologically incorrect or unfolded protein response (UPR) enables neoplastic cells to resist TKI therapeutics, such as imatininb [35].

#### 3.3.3. Fatty acid biosynthesis

Certain fatty acid metabolic markers guide resistance towards imatinib [36].

#### 3.3.4. Cellular catabolism

This representative class majorly includes monocarboxylic acid biosynthesis and fatty acid metabolism processes. Activation fatty-acid oxidation (FAO) (which is compensatory to imatinib mediated suppression of BCR-ABL and glycolysis), contribute to glucose-independent cancer cell survival and thereby resistance to Imatinib therapy [37]. Therefore, patients with activated FAO may exhibit Imatinib resistance.

#### 3.3.5. Tumor necrosis factor (TNF) production

This representative class includes processes like tumor necrosis factor (TNF) superfamily cytokine productions. Experimental evidence showed that FOXO3a activation could trigger increased expression of tumor necrosis factor-related apoptosis-inducing ligand (TRAIL) which could contribute to overcoming imatinib resistance. In general, FOXO3a functions downstream of Bcr–Abl tyrosine kinase in its phosphorylated inactive form in CML, which is reversed by Imatinib activity bringing FOXO3a in its activated form which further increases production of TNF family proteins. Hence TNF superfamily proteins’ production is an indication of imatinib sensitivity [38,39].

#### 3.3.6. Sulfur amino acid transport

Major sulfur containing amino acids in cells are methionine, cysteine, homocysteine, and taurine. Sulphur containing amino acids have great implications in the tumor microenvironment. However, evidence linking sulfur amino acid transport to Imatinib response is yet to be validated.

#### 3.3.7. Others

This group has different biological processes, such as axon extension, neurofibrillary tangle assembly, endocannabinoid signaling pathway, amyloid-beta formation and cell migration by vascular endothelial growth factor signaling pathway.

### 3.4. Network analysis

As was mentioned earlier, it is essential to know the regulatory link between identified genes and drug’s MoA or drug’s direct targets. To address this, BN was created for responders and non-responders, followed by comparison between two networks (Fig. 5A-C). The network architecture for responder samples (Fig. 5A) differed significantly from that of non-responder samples (Fig. 5B). The comparative figure (Fig. 5C) showed new connection that appeared (indicated with blue dashed line), disappeared (red solid line) or remained unchanged (black line) in non-responders.

**Fig. 5.**
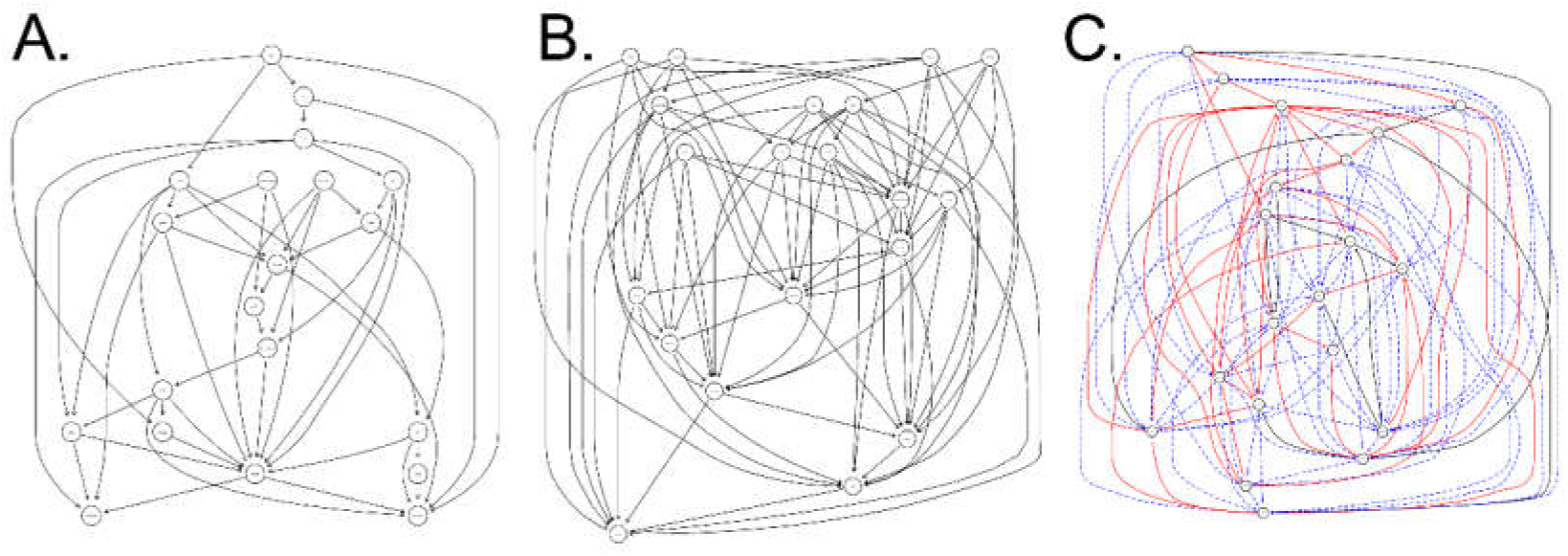
Bayesian network analysis. Bayesian network was created for (A) responders and (B) non-responders. (C) is the comparison between two networks. Blue dashed lines appear in non-responders while red lines disappear in them. Black lines remain unchanged in both cases.

BN analysis of the non-responder population identified one unique connection to them – ACADVL to PDIA5 (MIR7110) (Table 1). Both of these two proteins are well established for their roles in unfolded protein response (UPR) in the endoplasmic reticulum (ER). Recent studies suggested that these two proteins play important roles in cancer drug-resistance and closely interacts with GPR78 chaperon and transcription regulator TRF6α [40–43]. In our dataset also both ACADVL and PDIA5 were over-expressed in the non-responders group when compared to the expression patterns of the responders group. Further studies on the not so well understood UPR – ER Stress – Immune response mechanism could provide vital information on the development of novel therapeutics in several cancers [44].

Cortactin (CTTN) and Clusterin (CLU) are well established regulators of cancer metastasis and drug resistance [45–47] as well as are associated with the gene ontology biological process - negative regulation of apoptotic signaling pathway (GO:2001234). While CLU has a chaperone role, CTTN works as an actin cytoskeleton regulator downstream to Src kinase, another protein dis-regulated in different cancers [48]. While the direct relationship of CTTN and CLU are not well reported, over-expression of these two proteins is associated with poor cancer prognosis. In our dataset also, these two proteins exhibited over-expression in the non-responders group while responders exhibited lower expressions of both the proteins. CLU-CTTN connection being unique to the responders group (Table 4), it can be guessed that their lower expression is far more important for imatinib sensitivity than over expression mediated imatinib resistance.

**Table 4.** BN Comparative analysis. Comparative analysis of strong connections mapped from separate BN analysis of responding and non-responding population. While no common connections were identified across the non-responders and responders, seven unique connections (green text) were mapped to drug responding population compared to one unique connection (red text) coming from the non-responding group.

| From | To | Strength | Direction | Remark |
| --- | --- | --- | --- | --- |
| ABL1 | ACADVL | 0.88 | 0.76 | Connections present only in responding group |
| CSF1R | KIT | 0.8 | 0.81 |  |
| CSF1R | CTNS | 0.76 | 0.76 |  |
| CSF1R | CTSA | 0.82 | 0.78 |  |
| CSF1R | EMILIN1 | 0.86 | 0.8 |  |
| CLU | MEIS3P1 | 0.86 | 0.81 |  |
| CLU | CTTN | 0.99 | 0.9 |  |
| ACADVL | MIR7110.PDIA5 | 0.8 | 0.79 | Connections present only in non-responding group |

Putative homeobox protein Meis3-like 1 (MEIS3P1) protein is poorly understood but has been found to be dis-regulated in cancer related contexts [49–52]. Our data also suggest that its lower expression is potentially associated with better imatinib sensitivity. Similarly, ABL1 and ACADVL relationship under cancer context is not well established but both could potentially play important roles in SOCS3 mediated cancer prognosis. CSF1R is well known for having a beneficial effect on several cancers when inhibited by drugs [53] but its relation with BN predicted interactors like KIT, CTNS, CTSA and EMILIN1 are not yet well established. CTNS and CTSA are regulators of lysosome while EMILIN1 is known to have a positive impact on cancers [54–56]. Further studies would be required to decipher the interactions of these predicted interacting pairs.

BN helps to identify unique genetic or protein interactions through a completely data driven approach. Several gene connections identified in this study could be rationalized based on existing information from published articles while some were entirely new. While the individual roles of ER stress response proteins or lysosomal proteins are well established, their interplay with other cancer regulators is poorly understood. Few studies have highlighted the potential beneficial impact of these proteins or their in-situ dysregulation in several cancers. Further focused and consolidated experimental studies would be required to validate such findings as well to further understand the therapeutic potential of these genes when targeted by drugs.

## 4. Conclusion

In this work, supervised machine earning was employed on patients’ transcriptomics data and 12 potential biomarkers were identified. These biomarkers were found to be associated with certain biological phenomena, such as, regulation of autophagy, topologically incorrect proteins, fatty acid biosynthesis, TNF production etc, which have shown predominant effect regulating imatinib resistance. A deep dive with BN revealed exciting findings of 7 gene-gene causal network in responders and 1 gene-gene connection in non-responders that might play an instrumental role in drug response.

Interestingly, this method is disease or drug system independent and can be used in several kinds of biomarkers prediction and thereby stratification, either patient level or finding new indications/ disease subtype sensitive to therapy. In this current study, the aim was to identify drug response biomarkers for which drug linked response classes have been fed into the pipeline. However, if the aim is to find disease metastatic biomarkers then only benign and metastatic level gene information will enable the pipeline to the identification of metastatic regulating biomarkers. Moreover, Bayesian network itself is an effective way to find the causal regulatory network and thereby brings forth a mechanistic insight. Together this machine learning and bn will be very powerful in analyzing the drug response nature in light of personalized medicine.

In this work, a very small data set has been presented as an example case, yet the pipeline has identified significant genes which can be used as potential drug response biomarkers. The pipeline can be easily scaled up for a large set of data and is definite to produce relevant insight.

## 6. Acknowledgement

AD and SB want to thank Dr Saurabh Bundela for his guidance and assessment on this work.

## 7. About authors

SB and AD have equally contributed to this work, from hypothesis generation to experiment design and execution. SB and AD are senior scientific managers at Excelra knowledge solution, India, a contract research organization working in the pharma analytics. Both of them have Ph.D. in bioinformatics (2013-2016) and are trained in molecular biology techniques (2009-2013).

